# Histone H3K4 and H3K36 methylation promotes recruitment, but not activity, of the NuA3 histone acetyltransferase complex in *S. cerevisiae*

**DOI:** 10.1101/096511

**Authors:** Benjamin J.E. Martin, Kristina L. McBurney, Vicki E. Maltby, Kristoffer N. Jensen, Julie Brind’Amour, LeAnn J. Howe

## Abstract

Histone post-translational modifications (PTMs) alter chromatin structure by promoting the interaction of chromatin-modifying complexes with nucleosomes. The majority of chromatin-modifying complexes contain multiple domains that preferentially interact with modified histones, leading to speculation that these domains function in concert to target nucleosomes with distinct combinations of histone PTMs. In *S. cerevisiae*, the NuA3 histone acetyltransferase complex contains three domains, the PHD finger in Yng1, the PWWP domain in Pdp3, and the YEATS domain in Taf14, which *in vitro* bind to H3K4 methylation, H3K36 methylation, and acetylated and crotonylated H3K9 respectively. However the relative *in vivo* contributions of these histone PTMs in targeting NuA3 is unknown. Here we show that *in vivo* H4K4 and H3K36 methylation, but not acetylated or crotonylated H3K9, recruit NuA3 to transcribed genes. Through genome-wide colocalization and by mutational interrogation, we demonstrate that the PHD finger of Yng1, and the PWWP domain of Pdp3 independently target NuA3 to H3K4 and H3K36 methylated chromatin respectively. In contrast, we find no evidence to support the YEATS domain of Taf14 functioning in NuA3 recruitment. Collectively our results suggest that the presence of multiple histone-PTM binding domains within NuA3, rather than restricting it to nucleosomes containing distinct combinations of histone PTMs, can serve to increase the range of nucleosomes bound by the complex. Interestingly however, the simple presence of NuA3 is insufficient to ensure acetylation of the associated nucleosomes, suggesting a secondary level of acetylation regulation that does not involve control of HAT-nucleosome interactions

## Introduction

Eukaryotic DNA is packaged into a nucleoprotein structure known as chromatin, which consists of DNA, histones, and non-histone proteins. Histones are extensively post-translationally modified, with specific modifications reflecting activities occurring on the underlying DNA. For example, genes transcribed by RNA polymerase II (RNAPII) have acetylated and H3K4 tri-methylated histones (H3K4me3) at their 5’ ends and H3K36 tri-methylated histones (H3K36me3) over the gene body (Pokholok *et al.* 2005; Liu *et al.* 2005). In contrast, histone H2A glutamine 105 methylation and H2A.X serine 139 phosphorylation are associated with RNA polymerase I transcription and DNA double strand break repair respectively (Rogakou *et al.* 1998; Tessarz *et al.* 2014). It is generally accepted that these modifications promote the biological processes to which they are associated.

Although some histone post-translational modifications (PTMs) can directly alter chromatin structure, most function as recognition sites for histone PTM binding domains (Yun *et al.* 2011). Histone PTM binding domains are found in complexes that facilitate transcription or alter chromatin structure, such as basal transcription factors, chromatin-remodeling complexes and even enzymes that post-translationally modify histones. The majority of chromatin-modifying complexes have multiple histone PTM binding domains, the purpose of which has been the subject of much speculation. The generally weak affinity of these domains for their requisite histone PTM has led to the hypothesis that multiple histone-binding interactions are required to stabilize the binding of complexes to chromatin (Yun *et al.* 2011). Indeed, it is suggested that multiple histone-binding domains function synergistically to translate “a code” of histone PTMs into a single biological outcome (Strahl and Allis 2000). Alternatively, multiple histone binding domains could act independently to target a chromatin-binding complex to a range of genomic loci.

One complex containing multiple histone PTM binding domains is the NuA3 histone acetyltransferase (HAT) complex in *S. cerevisiae.* NuA3 contains six subunits: including the catalytic subunit Sas3, Nto1, Eaf6 and three histone PTM-binding proteins: Yng1, Pdp3, and Taf14 (John *et al.* 2000; Howe *et al.* 2002; Taverna *et al.* 2006). Yng1 contains a PHD finger which binds to H3K4 mono-,di- and trimethylation, with binding affinity increasing with the number of methyl groups. Pdp3 contains a PWWP domain, which recognizes H3K36me3. Taf14, through its YEATS domain, binds to acetylated (H3K9ac) and crotonylated (H3K9cr) histone H3K9. While the *in vitro* binding of these proteins to histone PTMs has been well characterized (Table S1), the relative contributions these histone PTMs make to NuA3 targeting *in vivo* remains unknown.

In mammalian cells, two HAT complexes appear analogous to NuA3: the MOZ/MORF complex and the HBO1-BRPF1 complex. Both complexes contain BRPF1 (bromodomain PHD finger protein 1) or one of its paralogs, BRPF2 or 3 (Doyon *et al.* 2006; Lalonde *et al.* 2013). BRPF1/2/3 share sequence similarity with yNto1, but additionally contain carboxy-terminal, H3K36me3-specific PWWP domains (Vezzoli *et al.* 2010), and thus may serve the role of both yNto1 and yPdp3 in mammalian complexes. The MOZ/MORF complex also has an H3K4me2/3- specific PHD finger in its subunit, ING5 (inhibitor of growth 5), while the HBO1–BRPF1 uses ING4 or ING5 to bind to H3K4me2/3 (Doyon *et al.* 2006; Lalonde *et al.* 2013). Finally, although the MOZ/MORF and HBO1-BRPF1 complexes lack a Taf14 equivalent, a bromodomain within BRPF1/2/3 and a double PHD finger in MOZ/MORF show specificity for acetylated histones (Laue *et al.* 2008; Qiu *et al.* 2012; Liu *et al.* 2012; Ali *et al.* 2012). Thus, the histone PTM binding domains in NuA3 are conserved in analogous complexes in other organisms, although the relative contribution that these domains make in targeting NuA3 has yet to be determined.

In this study we show that NuA3 is primarily targeted to mid-gene regions via interactions with H3K36me3 and H3K4me1/2/3, while H3K9ac and H3K9cr are unlikely to play a role in recruitment. Simultaneous disruption of H3K4 and H3K36 methylation abolishes NuA3 recruitment to actively transcribed genes. In contrast, disruption of H3K4 or H3K36 methylation singularly results in partial loss of NuA3 recruitment, suggesting that these PTMs recruit NuA3 independently and arguing against a synergistic effect. Finally, we show that NuA3 occupancy does not dictate histone acetylation indicating that controlled targeting is not the only mechanism for regulation of NuA3 function.

## Materials and Methods

### Yeast strains, antibodies and plasmids

All strains used in this study were isogenic to S288C, and are listed in Table S2. All strains are available upon request. Yeast culture and genetic manipulations were performed using standard protocols. Genomic deletions were verified by PCR analysis and whole cell extracts were generated as previously described (Kushnirov 2000). The previously described *kanMX-GAL1pr-flo8-HIS3* strains (Cheung *et al.* 2008) were generous gifts of Fred Winston.

### Drug treatments

For H3K23ac ChIP-seq, *bar1*Δ cells were arrested in G1 by 2.5 hour treatment with 5 μM α-factor. Culture synchrony in G1 was confirmed by the appearance of “shmooing” cells, as seen under the microscope. TSA was added for 15 minutes at 25 μM, from 5 mM DMSO stock.

### Chromatin immunoprécipitation with sequencing (ChIP-seq)

The ChIP-seq protocol was based on that outlined in (Maltby *et al.* 2012). Following cell lysis, the chromatin pellets were re-suspended in 900 μl of NP-S buffer (0.5 mM spermidine, 1 mM β-ME, 0.075% NP-40, 50 mM NaCl, 10 mM Tris pH 7.4, 5 mM MgCl_2_, 1 mM CaCl2) and digested with 400 units of microccocal nuclease at 37°C for 10 minutes to obtain predominantly mono-nucleosomal DNA. Reactions were stopped with the addition of 10 mM EDTA and digested lysates were clarified by centrifugation at 9000g for 5 minutes. To extract insoluble chromatin, pellets were re-suspended in 300 μl of lysis buffer with 0.02% SDS, and sonicated in a Diagenode Bioruptor at high output for 30 seconds on/30 seconds off for a total of 4 minutes. Extracts were then re-clarified by centrifugation at 9000g for 10 minutes, and the supernatant pooled with the pre-existing extract. The buffer composition of the lysate was adjusted to that of the original lysate buffer, and 10% was set aside as input. The supernatant was pre-cleared by incubation with 100 μl magnetic protein-G Dynabeads (Life Technologies) for 1 hour at 4°C. Pre-cleared lysates were incubated with 5 μg of αHA (Roche, cat no. 12013819001), αH3K4me3 (Abcam, cat no. ab1012), or αH3K23ac (Active Motif, cat no. 39131) antibodies at 4°C overnight. Immune complexes were precipitated by incubation with 100 μl of magnetic protein-G Dynabeads for one hour at 4°C. Beads were washed and the immunoprecipitated DNA subjected to Illumina HiSeq 2000 paired- end sequencing as described (Maltby *et al.* 2012). Reads were aligned to the *S. cerevisiae genome* (Saccer3 genome assembly) using BWA (Li and Durbin 2010). Average gene profiles were generated using the sitepro tool in the CEAS genome package (http://liulab.dfci.harvard.edu/CEAS/) and plotted using R. Values past the polyadenylation sequence (Park *et al.* 2014) were excluded from the average calculation, and the fraction of genes included in the average calculation at any given distance from the TSS shown. The sequencing data generated for this paper will be made available through the Gene Expression Omnibus (GEO) database, www.ncbi.nlm.nih.gov/geo.

#### Published datasets

The datasets for the co-occurring nucleosomes and for H3K4me3 in the H3K36R mutant (Sadeh *et al.* 2016) were downloaded from the SRA study SRP078243. Histone methylation and acetylation datasets from (Weiner *et al.* 2015) were downloaded from SRA study SRP048526. H3K9ac datasets from (Bonnet *et al.* 2014) were downloaded from SRA study SRP033513. The H3K4me3 dataset from (Maltby *et al.* 2012) was from our previous study. The fastq files were mapped to saccer3 using BWA (Li and Durbin 2010). The methylation data from (Schulze *et al.* 2011) was downloaded from www.yeastgenome.org as mapped MAT scores. The Yng1 data from (Taverna *et al.* 2006) was kindly provided by the authors as mapped intensity scores for IPs and inputs. The NET-seq data from (Churchman and Weissman 2011) was downloaded from SRA study SRP004431 and mapped to saccer3 using bowtie (Langmead *et al.* 2009). The processed data for Gcn5 from (Xue-Franzén *et al.* 2013) was downloaded from the GEO depository, series GSE36600. For average gene profiles, the +1 nucleosome was called using BEDTools (Quinlan and Hall 2010) as the closest consensus nucleosome position (Brogaard *et al.* 2012) to the TSS (Brogaard *et al.* 2012). Cell cycle regulated genes were identified by (Eser *et al.* 2014) and were excluded from analysis comparing G1-arrested and asynchronous datasets.

#### Nucleosome enrichments

For each dataset the average coverage over genome wide called nucleosome positions (Weiner *et al.* 2015) was calculated. When available the IPs were normalized to input files. Spearman correlation matrix was calculated in R, considering all pairwise complete observations.

#### Gene peak enrichment

For each gene, 100 bp windows were constructed in 5 bp steps, from upstream edge of + 1 NCP to the end of the polyadenylation site to a maxium of 3000 bp. The average signal (IP coverage for data from (Sadeh *et al.* 2016) and IP/input for all others) was calculated for each bin. The peak bin for each gene was calculated and location of peak enrichment. Boxplots for peak position were constructed for each of the indicated data tracks.

#### Boxplots

Méthylation enrichment defined as having an IP/input of greater than 1.5, and depletion corresponded to an IP/input of less than 0.75. Boxplots extend from the 1^st^ to 3^rd^ quartiles, with whiskers extending to 1.5 times the interquartile range or to the extreme of the data. Notches extend +/- 1/58 IQR /sqrt(n) and give an approximation of the 95% confidence interval for the difference in 2 medians.

### Chromatin immunoprécipitation with quantitative PCR (ChIP-QPCR) of Sas3

Cells were grown in 50 ml YPD to an OD600 of 0.8 and cross-linked with 1% formaldehyde for 30 minutes at room temperature. The reaction was stopped by the addition of 125 mM glycine, and incubated at room temperature for a further 15 minutes. Pellets were washed twice with cold PBS, resuspended in 600 μl; of lysis buffer (50 mM HEPES pH 7.5, 140 mM NaCl, 0.5 mM EDTA, 1% triton X-100, 0.1% sodium deoxycholate), and lysed mechanically by vortexing with glass beads for 25 minutes at 4°C. The lysates were spun down by centrifugation at 15,000g for 30 minutes, the supernatant was discarded and the pellet was washed and resuspended in 500 μL lysis buffer. The resuspended pellets were sonicated at high output for 30 seconds on/30 seconds off for 30 minutes to obtain an average fragment length of 250 bp. A further 200 μl of lysis buffer was added to each sample, and the lysates were clarified by centrifugation at 9,000g for 10 minutes. Ten percent of the lysate was reserved for input. Lysates were incubated with 1 μg antiHA antibody (Roche, cat no. 12CA5) at 4°C overnight, followed by precipitation of immune complexes with 25 μl protein G Dynabeads at 4°C for 1 hour. The DNA was eluted and processed as described previously (Maltby et al., 2012a) and qPCR was performed in technical triplicate using the primers listed in Table S3.

## Results

### NuA3 is primarily bound to mid-gene regions of actively transcribed genes

The NuA3 histone acetyltransferase complex contains three histone PTM binding domains: the Yng1 PHD finger, the Pdp3 PWWP domain, and the Taf14 YEATS domain, which show specificity for H3K4me1/2/3, H3K36me3 and H3K9ac/H3K9cr respectively (Table S1). To determine the relative contribution of each histone PTM in targeting the NuA3 complex, we reasoned that histone PTMs that promote the interaction of NuA3 with chromatin would co-localize with Sas3. To this end, we mapped NuA3-bound nucleosomes at high resolution *in vivo* using an MNase-based ChIP-seq approach, which has previously been used to map chromatin remodelers (Koerber *et al.* 2009; Floer *et al.* 2010; Yen *et al.* 2012; Ramachandran *et al.* 2015). We immunoprecipitated HA-tagged Sas3, in parallel with an untagged control, from crosslinked MNase-digested chromatin and performed paired-end sequencing. We could not detect any DNA in the untagged control mock IP, but nonetheless constructed a library and included the sample in the pool for sequencing. While sequencing of the Sas3 IP and inputs produced over 8 million DNA fragments, only 84,973 DNA fragments were recovered in the untagged control IP, confirming the specificity of our Sas3 ChIP-seq experiment.

We mapped the sequenced DNA fragments to the *S. cerevisiae* genome, aligned genes by the +1 nucleosome, and calculated the average profile for each sample (Figure 1A). Relative to the input Sas3 was enriched in the gene body, but not at the +1 nucleosome, and this enrichment was specific to the Sas3 IP and was not observed for the untagged control. We next compared Sas3 occupancy to a previously reported sonication-based mapping of Yng1 (Taverna *et al.* 2006). Unsurprisingly, Yng1 and Sas3 levels correlated positively genome-wide with a Spearman correlation coefficient of 0.48 (Figures 1B and S1). Additionally both of the Yng1 and Sas3 input-normalized distributions peaked in enrichment 600 bp to 800 bp downstream of the +1 nucleosome (Figure 1C). Thus our mapping of Sas3 agreed well with the previous mapping of Yng1, but the MNase-based approach increased the resolution of the assay, and we found that NuA3 was primarily recruited to ‘mid-gene’ regions containing the +5 to +7 NCPs (Figure 1C). We also observed that Sas3 was enriched on long genes and modestly enriched on genes transcribed by RNAPII (Figure 1D, E, and S2), consistent with Sas3 recruitment to mid-gene regions of actively transcribed genes.

**Figure 1:**
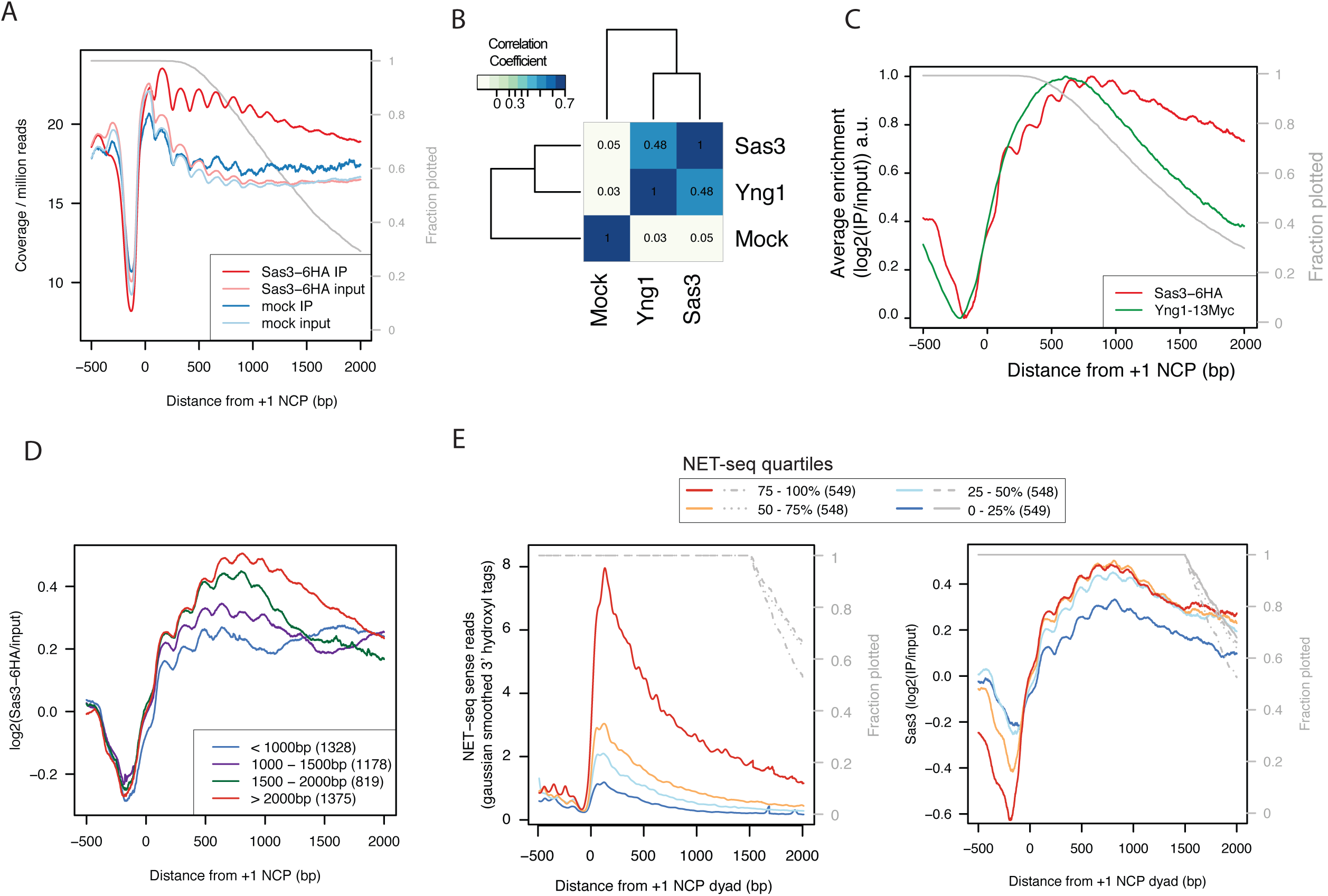
NuA3 is primarily bound to mid-gene regions of actively transcribed genes. A. The average sequence coverage relative to 4701 +1 nucleosomes for Sas3-HA and untagged IPs and MNase inputs. B. Spearman correlation coefficients for input-normalized Sas3 and Yng1 genome-wide enrichments. C. Average enrichment for Yng1 and Sas3 relative to the +1 nucleosome. Signal represents the log2(IP/input) and plotted as relative signal within each sample. D. Sas3 enrichment by gene length, with gene length defined as the distance from the +1 nuclesome to the polyadenylation site. E. Sas3 enrichment by quartiles of sense NET-seq (Churchman and Weissman 2011) signal for genes longer than 1500bp. Only genes longer than 1500bp were plotted to avoid gene length effects but results are similar when all genes analyzed (Figure S2). Except for gene length plot, all average plots only include data until the PAS, and the grey line represents the fraction of genes still being plotted.

### NuA3 is associated with H3K4me3, H3K4me2, H3K4me1, and H3K36me3 nucleosomes genome wide

As domains within NuA3 have been reported to bind to H3K4me3, H3K4me2, H3K4me1, H3K36me3, H3K9ac, and H3K9cr *in vitro* (Table S1), we hypothesized that if these modifications recruit NuA3 to nucleosomes *in vivo* they will correlate positively with Sas3. To conduct this meta-analysis, we made use of published genome-wide studies of these histone PTMs to calculate nucleosome-based Spearman correlations with Sas3 and Yng1 (Figure 2A, Table S4). Since crotonylation has not been mapped in *S. cerevisiae,* but colocalizes with acetylation in mammalian cells (Sabari *et al.* 2015), we used H3K9ac as a proxy for H3K9cr. To our surprise, H3K9ac did not correlate with Sas3 or Yng1, which suggests that despite Taf14 binding *in vitro,* it does not function in NuA3 recruitment *in vivo.* Conversely, H3K36me3, H3K4me3, H3K4me2 and H3K4me1 all correlated positively with Sas3 and Yng1, supporting their role in NuA3-nucleosome binding *in vivo.* Furthermore co-occurring H3K4me3 and H3K36me3 nucleosomes, identified through sequential IPs (Sadeh *et al.* 2016), strongly correlated with Sas3 with Spearman correlation coefficients above 0.5 (Figure 2A, Table S4). The dual H3K4me3 and H3K36me3 methylated nucleosomes also specifically grouped with Sas3 and Yng1 following Hierarchical clustering, consistent with NuA3 binding to nucleosomes through the combined effects of H3K36me3 and H3K4 methylation.

**Figure 2:**
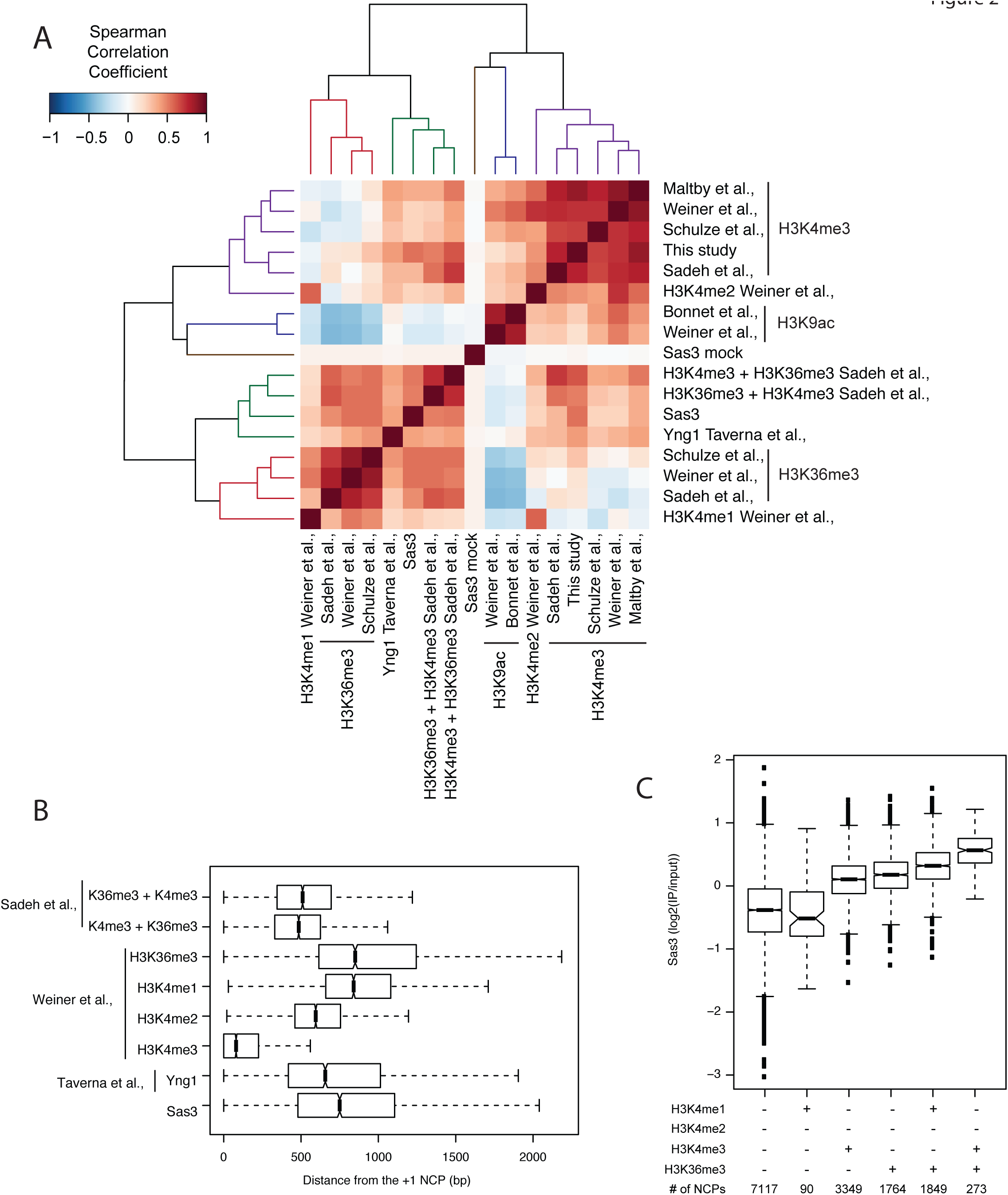
NuA3 is associated with H3K4me3, H3K4me2, H3K4me1, and H3K36me3 nucleosomes genome wide. A. Spearman correlation matrix for Sas3, Yng1 and histone PTM enrichments at 66360 genome-wide nucleosome positions. The rows and columns were sorted by hierarchical clustering, and the clustering is represented by dendrogram. B. Gene peak enrichments relative to the +1 nucleosome dyad represented by boxplot for 4701 genes. C. Sas3 enrichment at nucleosome positioned enriched or depleted for H3K4me1/2/3 and H3K36me3.

Next, we compared the locations of peak enrichment for Sas3, Yng1, H3K36me3, and H3K4 methylation (Figure 2B). Sas3 and Yng1 had median peak enrichments at 750 bp and 657 bp downstream of the +1 dyad respectively, which was in between the peak enrichments for H3K4me3 and H3K36me3. Notably Yng1 and Sas3 peaked in enrichment slightly downstream of co-occurring H3K4me3 and H3K36me3, suggesting that H3K4me2 and H3K4me1 are also important for NuA3 targeting. Altogether the peak enrichments of Sas3 and Yng1 were consistent with NuA3 being primarily recruited to mid-gene regions through the combined effects of H3K36me3 and H3K4 mono, di, and trimethylation.

We next assessed Sas3 enrichment on nucleosomes containing H3K4me1/2/3 or H3K36me3 singularly or in combination (Figure 2C). In the absence of H3K36me3 or H3K4 methylation Sas3 was depleted from nucleosomes. Likewise nucleosomes solely enriched in H3K4me1 were also depleted in Sas3, albeit with the caveat that we only identified 90 such nucleosomes and these may represent noise within one of the datasets or be atypical cases in the genome. H3K4me2 enriched nucleosomes are in all but a handful of nucleosomes also enriched for H3K4me1 or H3K4me3, and we were unable to interrogate this modification in isolation. Nucleosomes enriched singularly with H3K36me3 or H3K4me3 were enriched for Sas3 to a similar extent, suggesting that each of these modifications recruit Sas3 to nucleosomes independently. Nucleosomes enriched with both H3K4me1 and H3K36me3 were enriched for Sas3 to a greater extent than H3K36me3 alone, suggesting that H3K4me1 does indeed function in Sas3 recruitment. Nucleosomes enriched for both H3K4me3 and H3K36me3 were enriched further still in Sas3 binding, consistent with NuA3 preferentially binding H3K4me3 over H3K4me1. We found similar results upon analysis of Yng1 (Figure S3). Altogether this analysis supports H3K36me3 and H3K4me1/2/3 functioning to recruit NuA3 to chromatin *in vivo*.

### H3K4 and H3K36 methylation are necessary for Sas3 binding to active genes through Pdp3 and Yngl respectively

The genome-wide analysis suggested that H3K4 and H3K36 methylation are the major mechanisms for recruiting NuA3 to chromatin, and we next sought to test this directly. H3K4 and H3K36 methylation are entirely dependent on the histone methyltransferases Set1 and Set2 respectively, so we performed ChIP-qPCR for Sas3 in strains lacking the methyltransferases singularly and in combination. Mutation of either *SET1* or *SET2* individually resulted in a partial loss of Sas3 at *LOS1*, *SEC15*, *NUP145*, and *RPS28A* (Figure 3A). However Sas3 remained enriched over background at these four genes, and was also enriched relative to a repressed gene, *PUT4.* In contrast to the modest loss in the single mutants, the *set1Δ set2Δ* double mutant reduced Sas3 recruitment to background levels, demonstrating that Set1 and Set2 are necessary for Sas3 recruitment to chromatin. Furthermore, the Set1- dependence of Sas3 binding to the mid and 3’ regions of *LOS1*, *SEC15*, and *NUP145* occurred in regions depleted for H3K4me3 (Figure 3B) but enriched for H3K4me2 and H3K4me1 (Figure S4), which supports these modifications recruiting Sas3 *in vivo.* Set2-dependent binding of Sas3 occurred at regions containing H3K36me3 (Figure 3C), consistent with this PTM recruiting Sas3 *in vivo*. Additionally, from the single to double mutants, Sas3 displayed a stepwise reduction in binding, consistent with H3K36me3 and H3K4me1/2/3 recruiting Sas3 to chromatin in an independent and additive manner.

**Figure 3:**
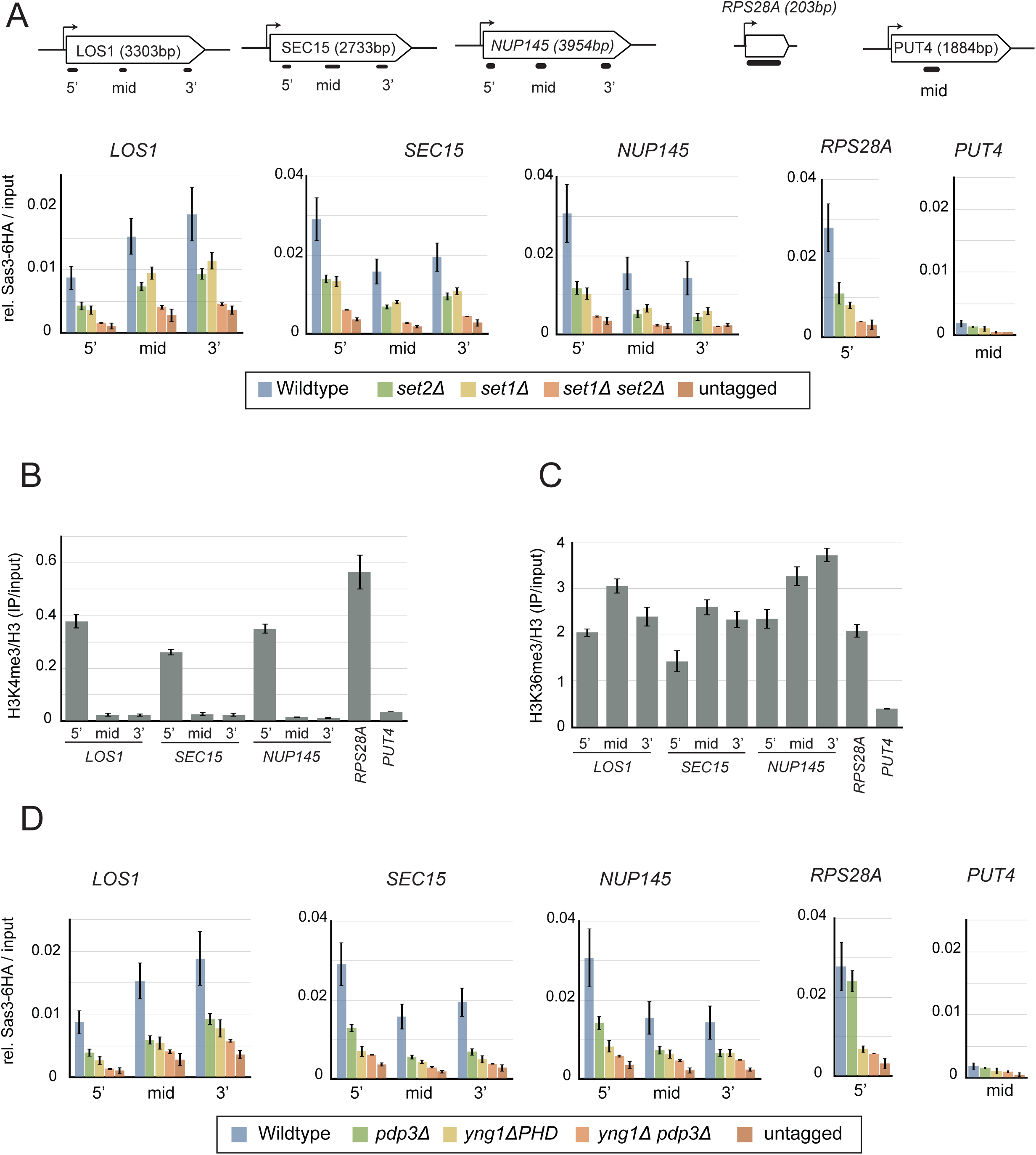
H3K4 and H3K36 methylation are necessary for Sas3 binding to active genes through Pdp3 and Yng1 respectively. A,C. Sas3 ChIP-qPCR in indicated strains at *LOS1*, *SEC15*, *NUP145*, *RPS28A*, and *PUT4*. Primer positions on genes are indicated in schematic. B. H3K4me3 and H3K36me3 ChIP-qPCR in wild type cells. Values represent the mean of at least three independent replicates. Error bars represent the standard error of the mean.

To test if Sas3 recruitment to chromatin was dependent on the NuA3 domains reported to bind histone methylation we performed ChIP-qPCR for Sas3 in *yng1ΔPHD*,*pdp3Δ*, and *yng1ΔPHD* strains (Figure 3D). The *pdp3Δ* mutant had a similar reduction in Sas3 binding as the *set2Δ* mutant at all but one locus, and demonstrates for the first time that Pdp3 is necessary for NuA3 recruitment to chromatin *in vivo.* At *RPS28A*, disrupting H3K36 methylation causes a reduction in H3K4me3 (Figure S5), so the greater loss of Sas3 binding in the *set2Δ* compared to the *pdp3Δ* mutant was likely an indirect effect. The*yng1ΔPHD* mutation caused a reduction in Sas3 recruitment at 5’, mid, and 3’ regions, consistent with Yng1 targeting NuA3 to H3K4me3, H3K4me2, and H3K4me1. The *yng1ΔPHD pdp3Δ* double mutant resulted in Sas3 recruitment comparable to the untagged control at all loci tested. Altogether our ChIP-qPCR results support the hypothesis that Sas3 binds to actively transcribed chromatin due to binding of H3K4me1/2/3 and H3K36me3 by the Yng1 PHD finger and the Pdp3 PWWP domain respectively.

### Sas3 occupancy does not dictate histone H3K23 acetylation

Histone H3K14 and H3K23 acetylation localize to the 5’ ends of genes, which is inconsistent with our demonstrated occupancy of Sas3 (Fig S6) (Weiner *et al.* 2015). One explanation for this discrepancy is the presence of the histone deacetylase complex (HDAC), Rpd3S, which deacetylates nucleosomes in the body of transcribed genes (Carrozza *et al.* 2005; Keogh *et al.* 2005; Joshi and Struhl 2005). To test whether HDAC inhibition could reveal acetylation in the bodies of genes, we performed ChIP-seq analysis for H3K23ac in cells treated with the HDAC inhibitor, trichostatin A (TSA) (Figure 4A). Following TSA treatment, H3K23ac spread further downstream of the TSS, but this PTM was still largely restricted to the 5’ ends of genes contrary to NuA3 occupancy, which was found across the bodies of genes. Further, deletion of *SAS3* failed to suppress cryptic transcription initiation of a reporter construct in a strain lacking the TSA-sensitive HDAC, RPD3S (Figure 4B) (Cheung *et al.* 2008), again suggesting that Sas3 was not responsible for acetylation at downstream targets.

**Figure 4:**
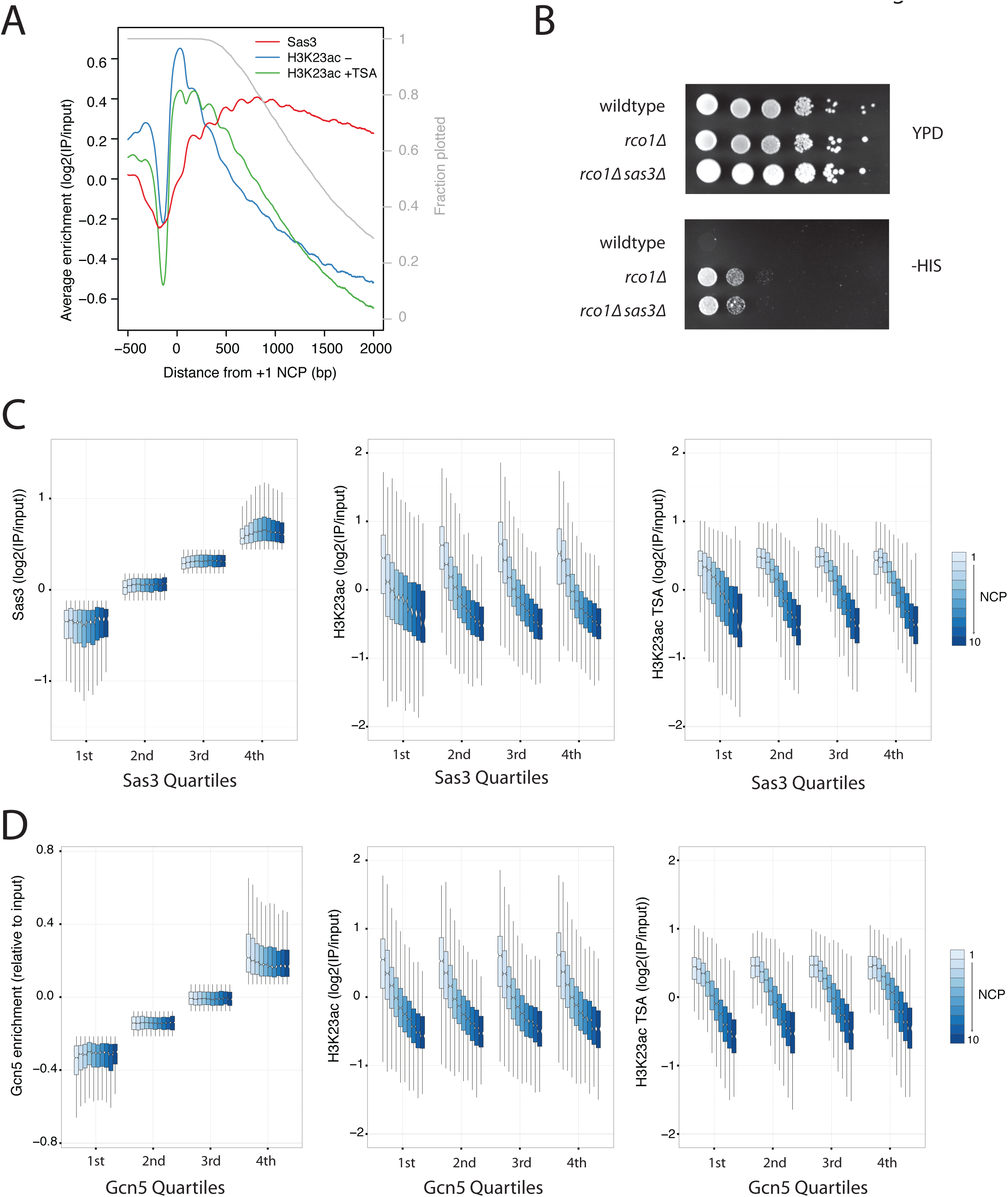
Sas3 occupancy does not dictate histone H3K23 acetylation. A. The average enrichment relative to 4264 +1 nucleosomes for Sas3 and H3K23ac before and after 15 minute incubation with 25μM TSA. Cell cycle regulated genes were excluded from this plot. Each gene is only included in the average calculation until its polyadenylation signal. The fractions of genes still contributing to the average profile are represented by the gray line. B. Ten-fold serial dilutions of the indicated strains containing the *kanMX-GAL1pr-flo8-HIS3* reporter were plated on rich media (YPD) and complete synthetic media lacking histidine, and incubated at 30°C for four days. C,D. HAT and H3K23ac before and after TSA enrichments by nucleosome position and by Sas3 (C) or Gcn5 (Xue-Franzén *et al.* 2013) (D) quartiles, represented as boxplots. Nucleosomes from cell-cycle regulated genes were excluded, leaving 33942 nucleosomes from +1 to +10 positions relative to the TSS. Outliers were not plotted

A disconnect between Sas3 occupancy and H3K23ac levels is surprising as it is generally thought that histone acetylation is regulated through control of HAT targeting. To further confirm these observations, we assessed the relationship between Sas3 and H3K23ac at the +1 to +10 nucleosomes, and while we did observe a modest association between Sas3 occupancy and H3K23ac (Figure S7A), the predominant determinant of H3K23ac was the nucleosome position relative to the TSS (Fig 4C). H3K23ac decreased into the gene body, and this was largely independent of Sas3 occupancy and was seen with or without TSA treatment. The H3 HAT, Gcn5, with Sas3, is necessary for global H3 acetylation (Maltby *et al.* 2012), and so we asked if Gcn5 occupancy (Xue-Franzén *et al.* 2013) could explain the disparity between Sas3 occupancy and H3K23ac. Similar to Sas3, Gcn5 displayed a subtle association with H3K23ac (Fig S7B), but again this was a modest effect compared to gene position (Figure 4D). We observed similar effects when selecting for nucleosomes enriched for one or both of Sas3 and Gcn5 (Fig S8). Thus the lack of association of Sas3 with H3K23ac cannot be explained by Gcn5 occupancy, and the similar lack of association of Gcn5 with H3K23ac suggests that regulating HAT activity post-recruitment may be a general phenomenon. Collectively these results indicate that while histone methylation promotes the association of Sas3 with chromatin in gene bodies, this does not necessarily result in histone acetylation.

## Discussion

In this study we investigated the relative contributions of histone PTMs in targeting NuA3 to chromatin. Using both genome-wide and locus-specific approaches we showed that H3K36me3 and H3K4me1/2/3 both independently and additively promoted the association of Sas3 with chromatin. We provide the first *in vivo* evidence for Pdp3 recruiting NuA3 to H3K36 trimethylated chromatin and for Yng1 recruiting NuA3 to H3K4me2 and H3K4me1. The additive effects of H3K36me3 and H3K4me1/2/3 resulted in NuA3 being primarily recruited to the +5 to +7 nucleosomes, approximately 700 bp into the gene. This is close to but slightly further into the gene body than where the mammalian homologue MOZ/MORF is predominantly found, approximately 400 bp downstream of the TSS (Lalonde *et al.* 2013). This 5’ shift in MOZ/MORF localization could be due to the mammalian complex’s reduced affinity for H3K4me1 (Champagne *et al.* 2008) (Table S1) and H3K36me3 (Vezzoli *et al.* 2010; Wu *et al.* 2011) (Table S1), resulting in a much greater dependence on H3K4me2/3 for recruitment to chromatin. The differing methyl-histone binding properties of NuA3 and MOZ/MORF coincide with differing genome-wide localization of H3K4me1 and H3K36me3. With longer mammalian genes H3K36me3 can stretch more than 20 kb from the TSS, while H3K4me1 is associated with enhancers (Barski *et al.* 2007). Thus the reduction in binding affinity for H3K36me3 and H3K4me1 has maintained MOZ/MORF targeting close to but downstream of the TSS, which suggests a conserved function in this genomic region.

Unlike for histone methylation, we found no evidence to support a role for histone acetylation or crontylation in NuA3 recruitment. While we did not test the effect of loss of acetylation or Taf14 on NuA3 binding, the complete loss of Sas3 recruitment in the absence of H3K4 and H3K36 methylation demonstrated that the YEATS domain was not sufficient for NuA3 recruitment. While it is possible that this domain has no function in NuA3, the retention of acetyl-binding domains in mammalian MOZ/MORF argues for a functional role. It is possible that binding to acetyl or crontyl lysines of the histone tail may regulate NuA3’s activity, similar to the role of histone methylation in stimulating the activity of the Rpd3S deacetylase complex (Govind *et al.* 2010). Alternatively regulation could occur through binding to non-histone substrates. Indeed proteomic studies show that Nto1 and Taf14 both contain acetylated lysines (Henriksen *et al.* 2012), and so it is possible that the YEATS domain is binding to a modified lysine in the NuA3 complex. Such a role is seen for the Rsc4 bromodomain, which binds to an acetylated lysine in the complex to regulate its function (VanDemark *et al.* 2007; Choi *et al.* 2008).

Our results showed that Sas3 was localized across the body of transcribed genes, which is inconsistent with the predominantly TSS-proximal patterns of H3K14ac and K23ac (Weiner *et al.* 2015). This is unsurprising however as other studies have shown that HAT occupancy is a poor predictor of histone acetylation (Xue-Franzén *et al.* 2010; Rossetto *et al.* 2014). Instead, these results suggest that there is a level of regulation of histone acetylation that is independent of HAT recruitment. Molecular simulation studies predict histone tails to be tightly intertwined with nucleosomal DNA (Shaytan *et al.* 2016), and thus disruption of these interactions may be required for acetylation by available HATs. Interestingly, although RNAPII is also found across gene bodies, NETseq and PAR-CLIP experiments show that RNAPII struggles to transcribe through the 5’ ends of genes (Churchman and Weissman 2011; Schaughency *et al.* 2014), where the majority of histone acetylation is found. Thus an attractive hypothesis is that histones are acetylated by available HATs in response to DNA unwrapping during slow RNAPII passage. Although other molecular mechanisms could explain our observations, our data underscores the fact that the presence of a histone PTM-binding domain within a chromatin-modifying complex does not ensure that the associated enzymatic activity will function on all nucleosomes with the requisite PTM.

## Acknowledgements

This work was supported by a Discovery Grant from the Natural Sciences and Engineering Research Council (NSERC) and an operating grant from the Canadian Institutes of Health Research awarded to L.J.H. B.J.E.M. is a recipient of an NSERC CGS award, while V.E.M. was supported by a University of British Columbia Four Year Fellowship. We gratefully acknowledge Fred Winston for providing yeast strains and plasmids and Sean Taverna for sharing ChIP-chip data.

---

**Figure S1:**
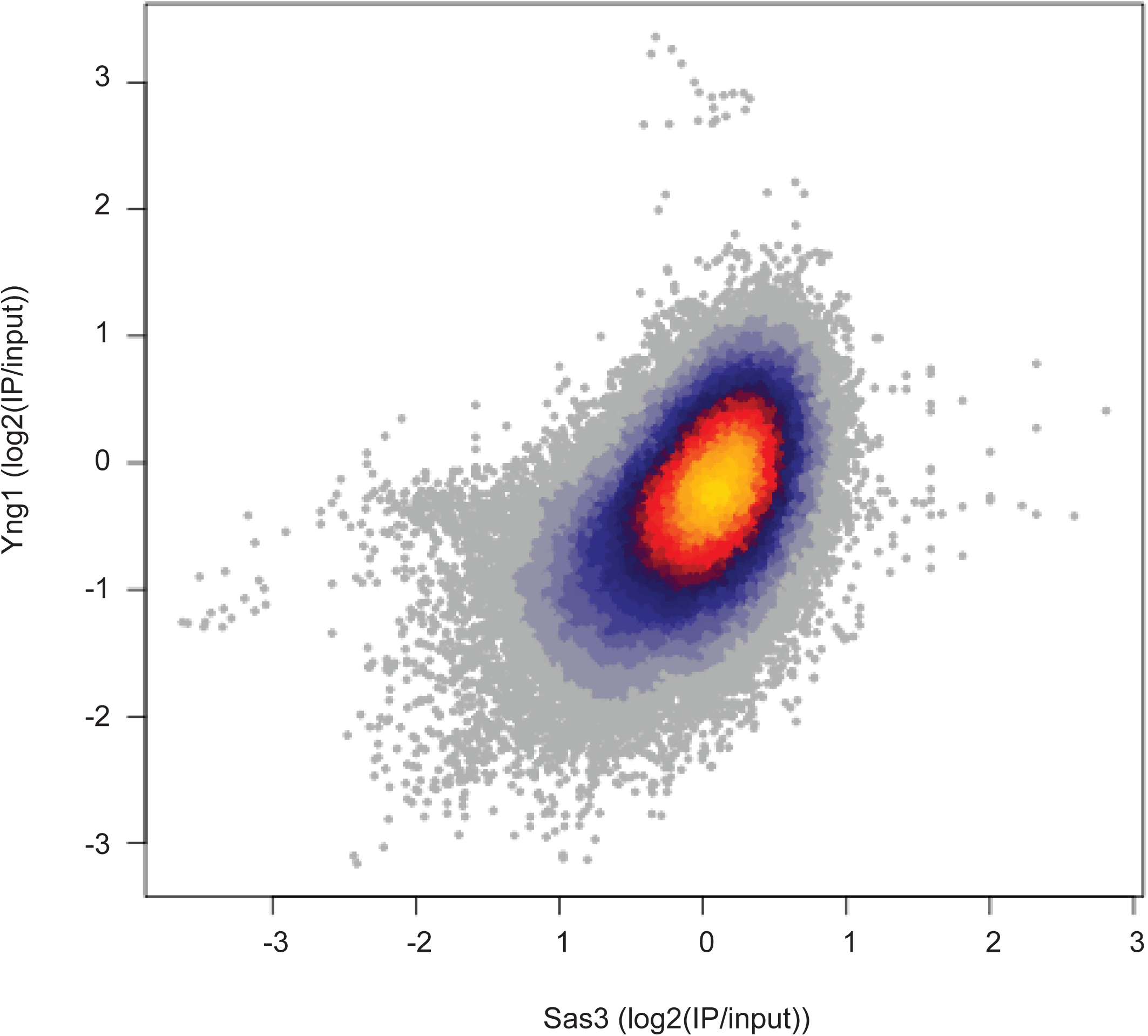
Sas3 and Yng1 colocalize genome wide. A. Scatter plot of Yng1 and Sas3 enrichment on 500 bp windows, stepping in 250 bp increments genome wide.

**Figure S2.**
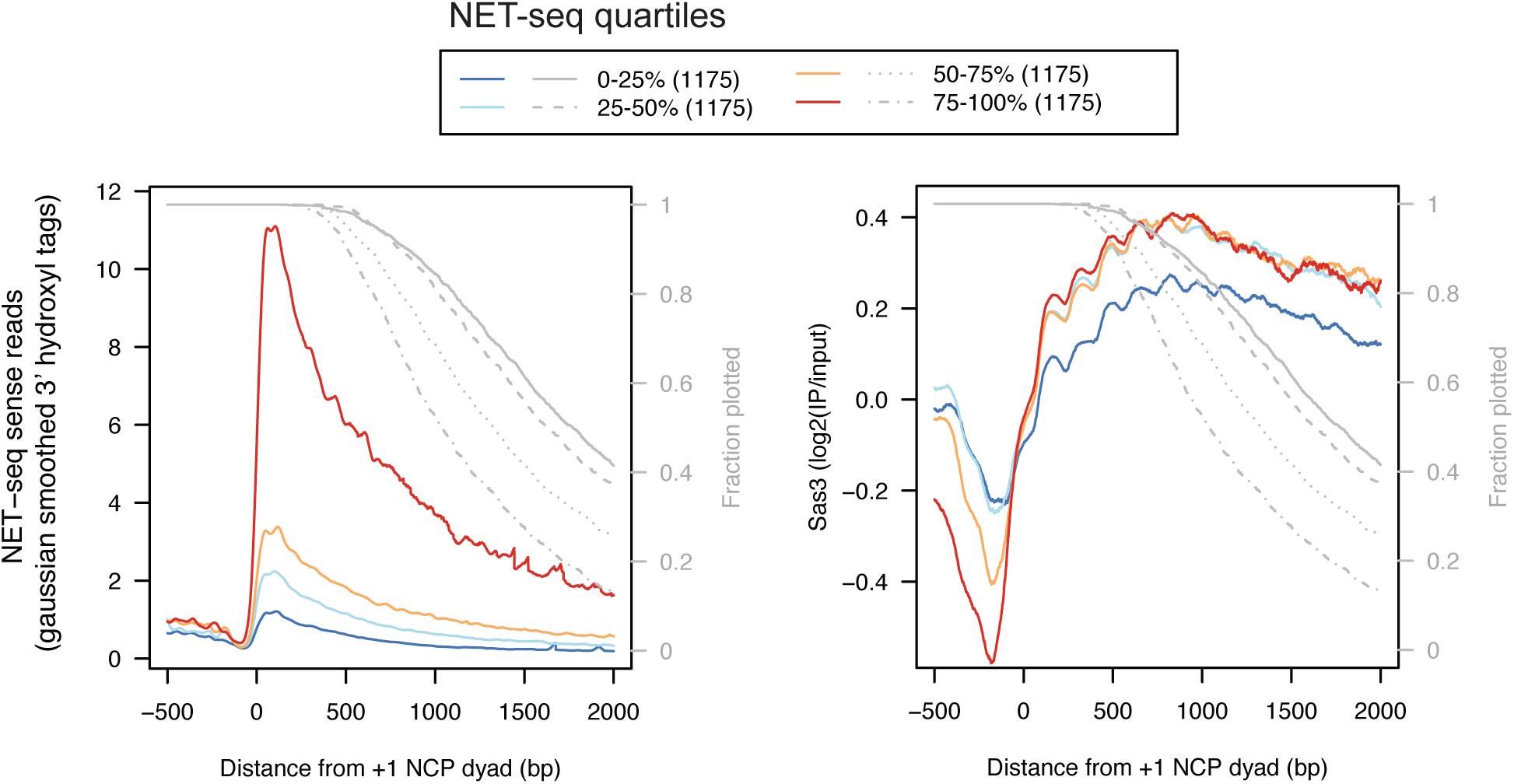
Sas3 is enriched on sense RNAPII-transcribed genes. A. Average profiles for NET-seq and Sas3 relative to the +1 nucleosome for genes split into quartiles of enrichment of NET-seq signal. Each gene is only included in the average calculation until its polyadenylation signal. The grey line represents the fraction of genes still contributing to the average profile.

**Figure S3:**
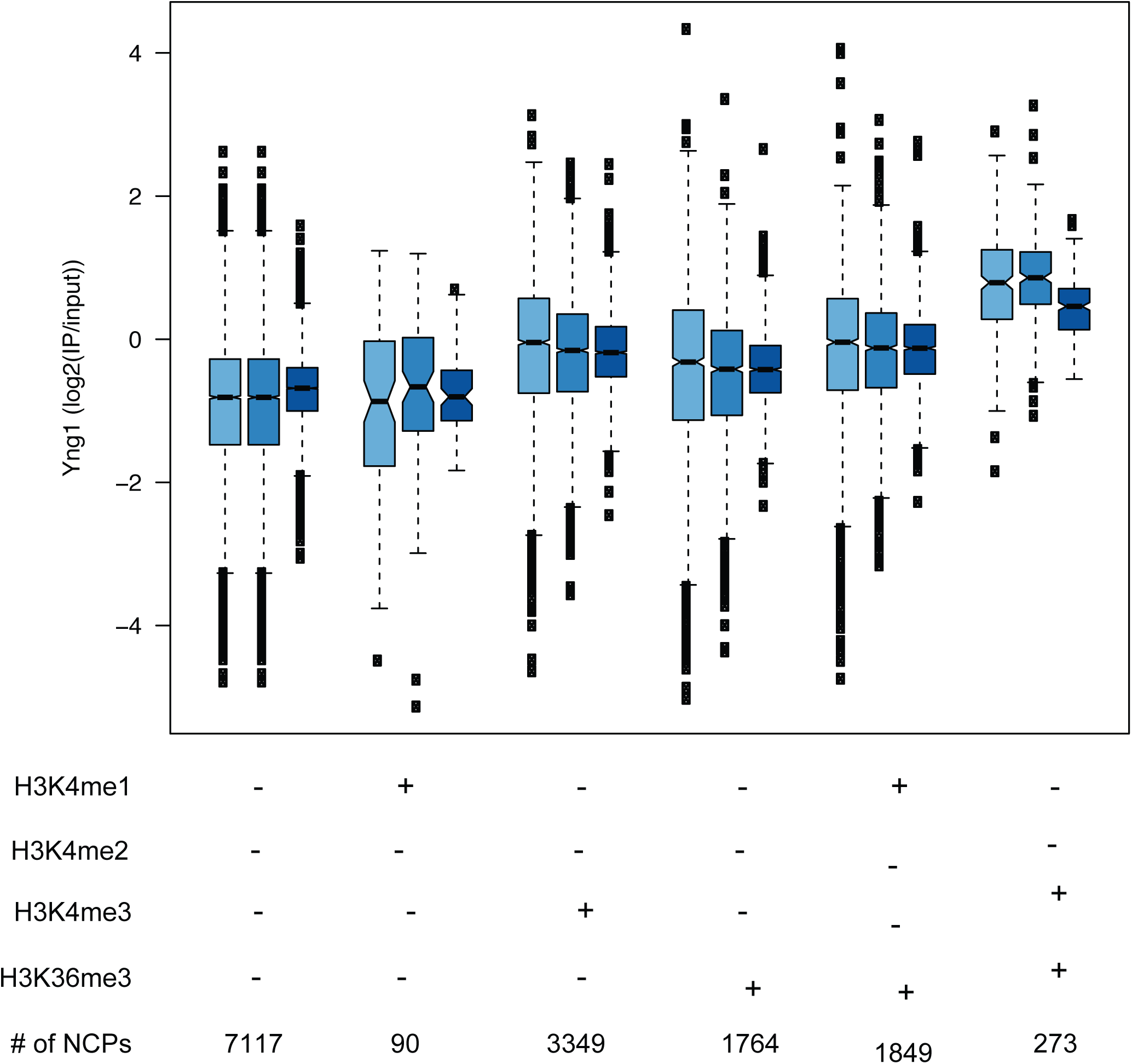
Yng1 is enrichment at methylated nucleosomes. Yng1 enrichment at nucleosome positioned enriched or depleted for H3K4me1/2/3 and H3K36me3. Yng1 replicate is indicated by colour.

**Figure S4:**
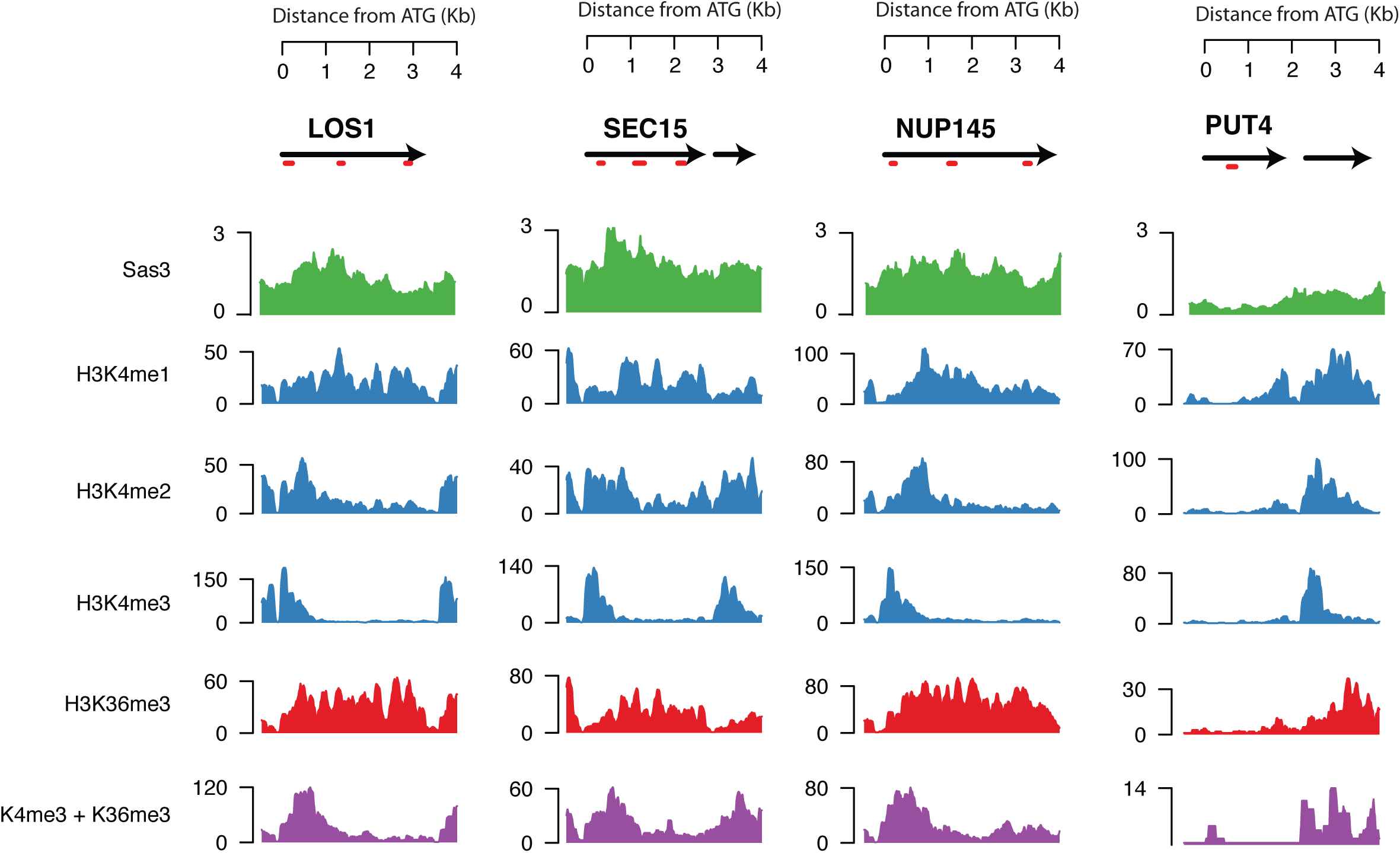
Histone methylation and Sas3 at candidate genes. Input-normalized Sas3 and histone methylation ChIP-seq coverage at *LOS1*, *SEC15*, *NUP145*, and *PUT4* genes. ChIP-qPCR amplicons are indicated by red bars on schematic.

**Figure S5:**
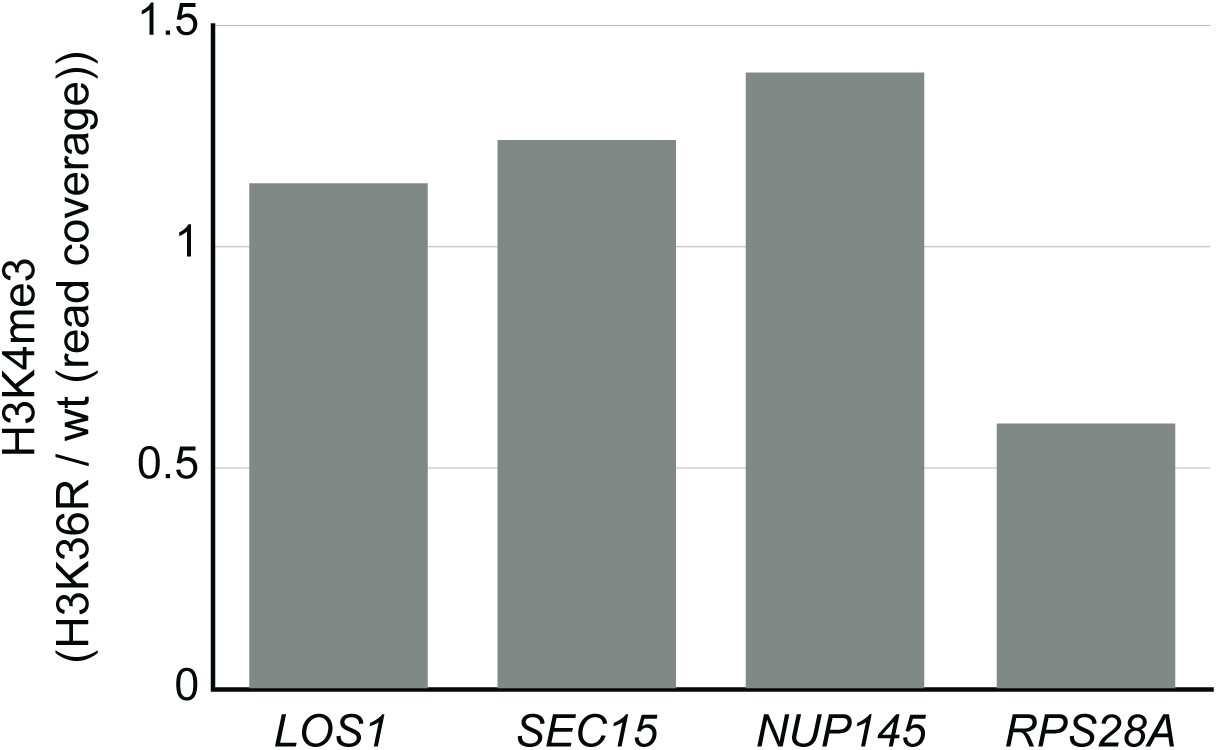
H3K4me3 decreases at *RPS28A* when H3K36 methylation is disrupted. H3K4me3 from wild type and H3K36R strains was quantitatively compared by competitive immunoprecipitation (data from Sadeh *et al.* 2016). The amount of H3K4me3 in wild type and H3K36R at 5’ qPCR loci (+100 bp on either side to account for change in resolution) was determined, and the change in methylation is plotted.

**Figure S6:**
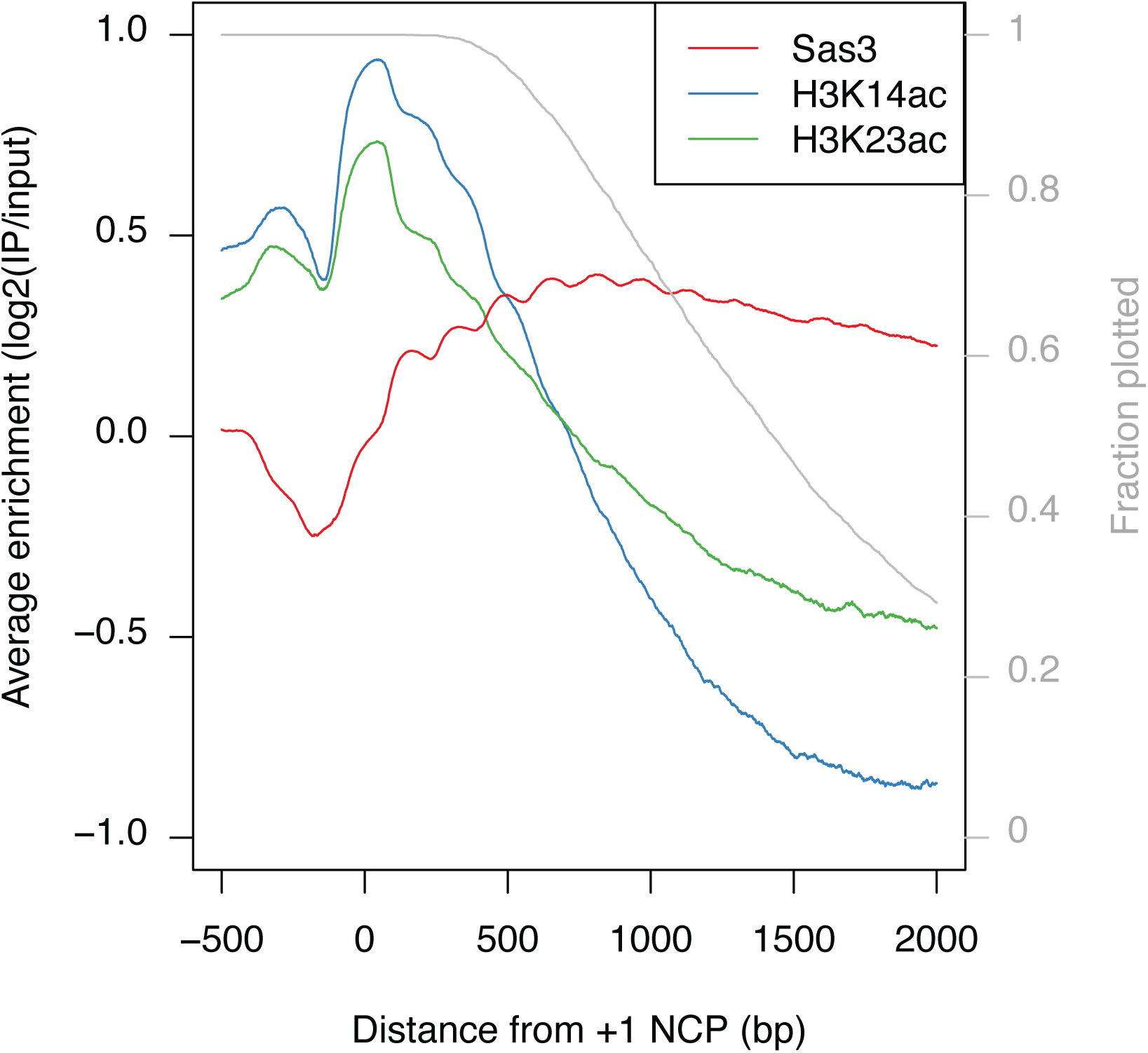
H3K14ac and H3K23ac are enriched 5’ of Sas3. The average enrichment relative to 4701 +1 nucleosomes for Sas3, H3K14ac, and H3K23ac. H3K14ac and H3K23ac data is from (Weiner *et al.* 2015). Each gene is only included in the average calculation until its polyadenylation signal. The fractions of genes still contributing to the average profile are represented by the gray line.

**Figure S7:**
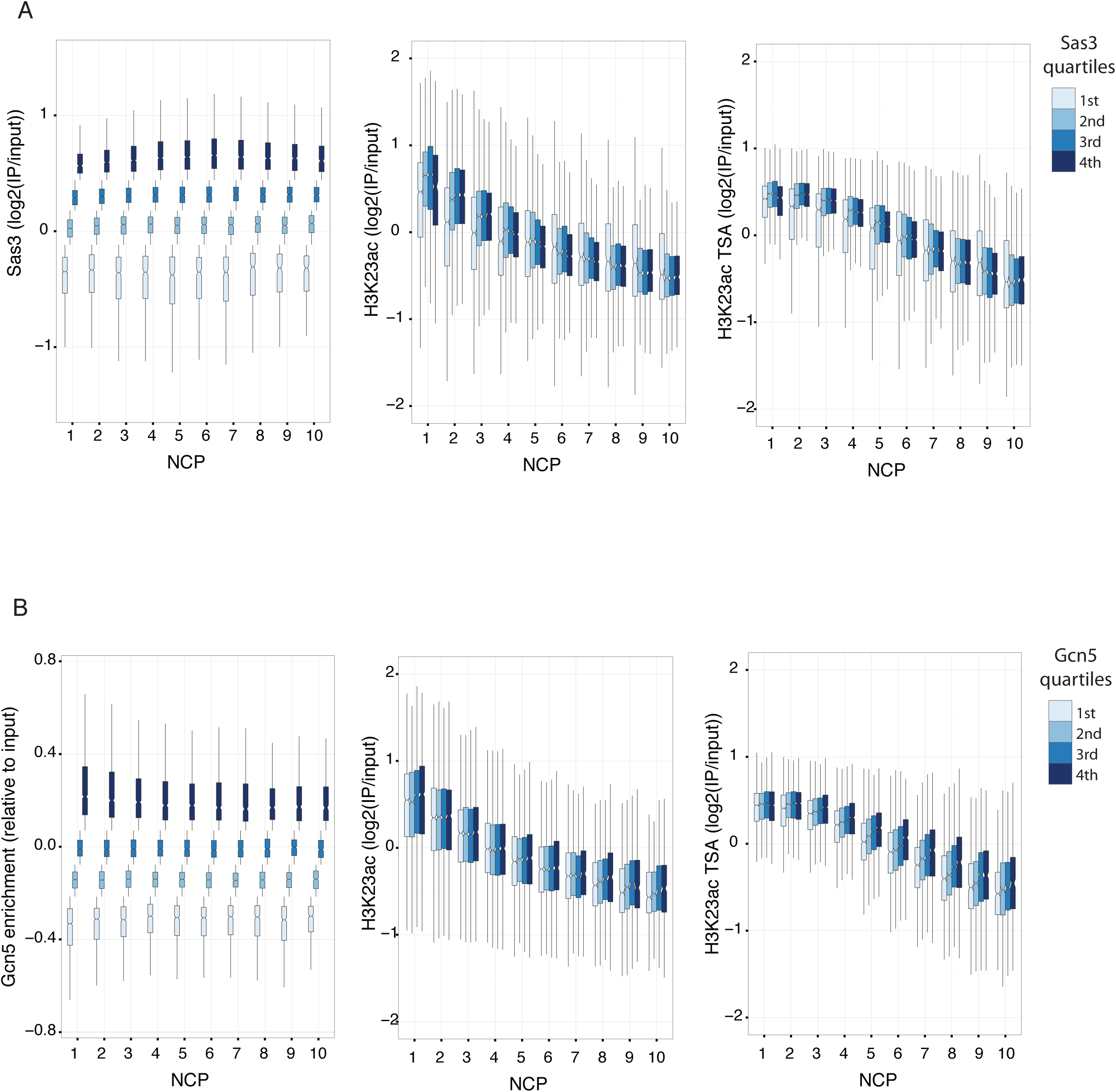
Modest association between Gcn5 or Sas3 and H3K23ac. HAT and H3K23ac before and after TSA enrichments by Sas3 (A) or Gcn5 (B) quartiles and by NCP, represented as boxplots. Nucleosomes from cell-cycle regulated genes were excluded, leaving 33942 nucleosomes from +1 to +10 positions relative to the TSS. Outliers were not plotted.

**Figure S8:**
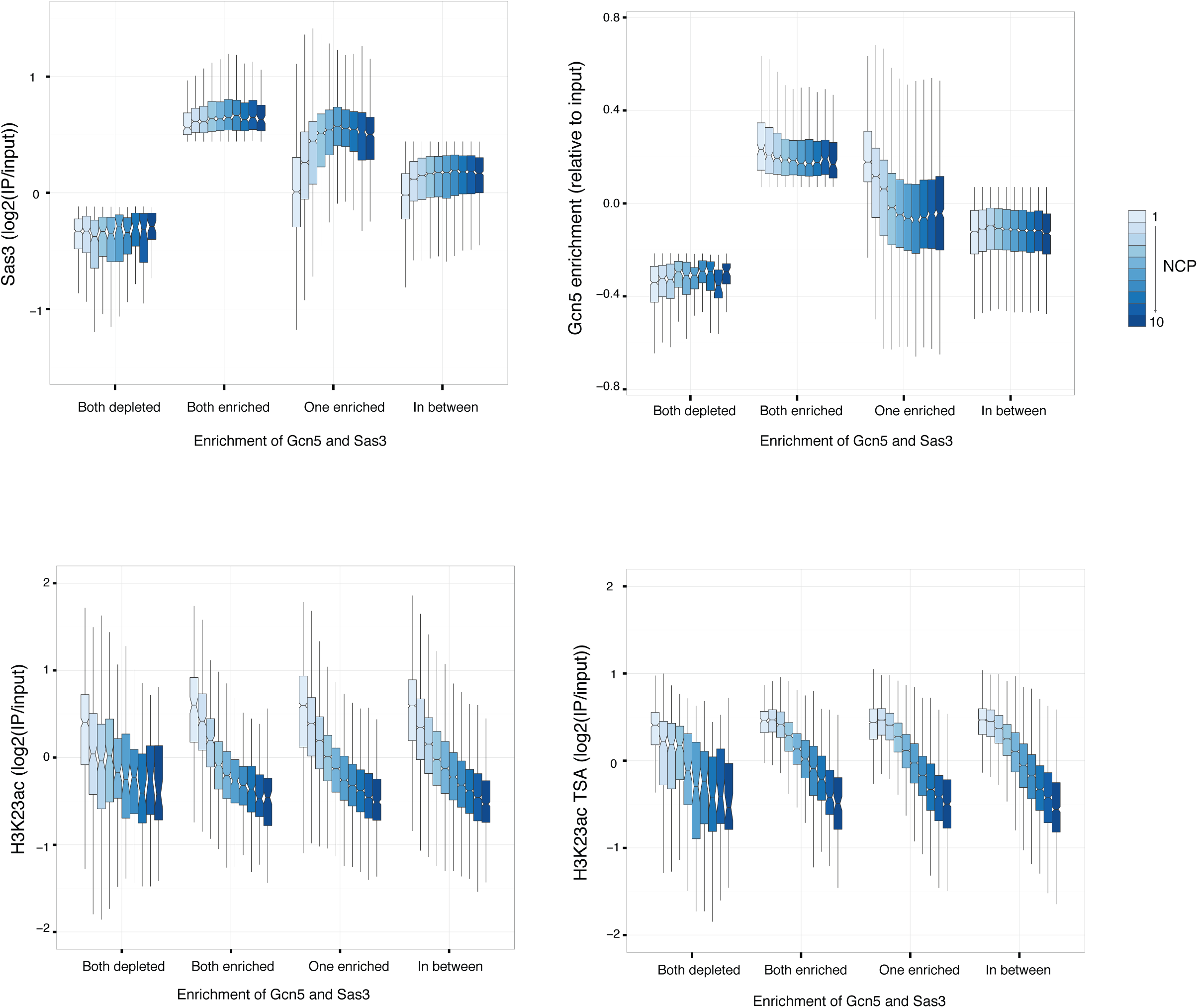
Gcn5 and Sas3 enrichment together does not dictate H3K23ac. Nucleosomes were classified as being enriched or depleted based on being the top or bottom quartile of HAT occupancy respectively. Nucleosomes were grouped as being depleted or enriched for both of Gcn5 or Sas3, enriched for one but not the other, and for not being in one of the above categories. Sas3, Gcn5, and H3K23ac before and after TSA treatment enrichments for each category were visualized by boxplot. Outliers were not plotted.

**Table S1:** Published *in vitro* dissociation constants for NuA3 and MOZ/MORF histone-PTM binding domains

| Domain | Peptide | $K_D$ (μM) | Reference |
| --- | --- | --- | --- |
| Yng1 PHD finger | H3K4me3 | 2.3 +/- 0.9,<br>9.1 +/- 1.6 | (Shi *et al.* 2007),<br>(Taverna *et al.* 2006) |
|  | H3K4me2 | 21.4 +/- 3.0 | (Taverna *et al.* 2006) |
|  | H3K4me1 | 50.7 +/- 7.6 | (Taverna *et al.* 2006) |
|  | H3K4me0 | >400 | (Taverna *et al.* 2006) |
| Pdp3 PWWP | H3K36me3 | 69.5 +/- 3.7 | (Gilbert *et al.* 2014) |
|  | H3K36me2 | 414 +/- 23 | (Gilbert *et al.* 2014) |
|  | H3K36me1 | >1000 | (Gilbert *et al.* 2014) |
|  | H3K36me0 | >1000 | (Gilbert *et al.* 2014) |
|  | H3K79me3 | 434 +/- 49 | (Gilbert *et al.* 2014) |
|  | H4K20me3 | > 1000 | (Gilbert *et al.* 2014) |
| Taf14 YEATS | H3K9ac | 150.6 +/- 14.5 | (Shanle *et al.* 2015) |
|  | H3K9,14,18ac | 49.9 +/- 2 | (Shanle *et al.* 2015) |
|  | H3K9cr | 9.5 +/- 0.5 | (Andrews *et al.* 2016) |
| ING5 PHD finger | H3K4me3 | 2.4 +/- 1.0 | (Champagne *et al.* 2008) |
|  | H3K4me2 | 16 +/- 1.2 | (Champagne *et al.* 2008) |
|  | H3K4me1 | 222 +/- 17 | (Champagne *et al.* 2008) |
|  | H3K4me0 | 261 +/- 34 | (Champagne *et al.* 2008) |
| BRPF1 PWWP | H3K36me3 | 2700 +/- 200,<br>2900 - 4000 | (Vezzoli *et al.* 2010),<br>(Wu *et al.* 2011) |
|  | H3K36me2 | very weak | (Wu *et al.* 2011) |
|  | H3K36me0 | NB | (Wu *et al.* 2011) |
|  | H3K4me3 | NB | (Wu *et al.* 2011) |
|  | H3K79me3 | very weak | (Wu *et al.* 2011) |
|  | H3K9me3 | NB | (Wu *et al.* 2011) |

**Table S2:** Strains

| Strain | Parent | Mating type | Genotype | Source |
| --- | --- | --- | --- | --- |
| YLH101 | FY602 | Mat a | *his3D200 leu2D1 lys2-128d ura3-52 trp1D63* |  |
| YVM138 | YLH101 | Mat a | *his3D200 leu2D1 lys2-128d ura3-52 trp1D63 SAS3-6HA::TRP* |  |
| YVM142 | YLR008 x YLH354 | Mat a | *his3D200 leu2D1 lys2-128d ura3-52 trp1D63 SAS3-6HA::HIS set1Δ::KANMX6 set2Δ::TRP* |  |
| YVM147 | YVM144 | Mat a | *his3D200 leu2D1 lys2-128d ura3-52 trp1D63 SAS3-6HA::TRP yng1ΔPHD::KANMX6* |  |
| YVM157 | YVM146 | Mat a | *his3D200 leu2D1 lys2-128d ura3-52 trp1D63 SAS3-6HA::TRP set1Δ::HISMx6* |  |
| YVM158 | YVM146 | Mat a | *his3D200 leu2D1 lys2-128d ura3-52 trp1D63 SAS3-6HA::TRP set2Δ::HISMx6* |  |
| YVM207 | YVM146 | Mat a | *his3D200 leu2D1 lys2-128d ura3-52 trp1D63 SAS3-6HA::TRP ylr455w::HIS* |  |
| YLH696 | YVM147 | Mat a | *his3D200 leu2D1 lys2-128d ura3-52 trp1D63 SAS3-6HA::TRP yng1DPHD::KANMX6 ylr455w::HISMx6* |  |
| YLH787 | YLH101 | Mat a | *his3D200 leu2D1 lys2-128d ura3-52 trp1D63 bar1::KAN* |  |
| FY2173 | | MATα | *his3Δ200 leu2Δ1 lys2-128Δ trp1Δ63 ura3-52 kanMX-GAL1pr-FLO8-HIS3* | (Cheung et al. 2008) |
| L1106 |  | Mat a | *his3Δ200 ura3Δ0 kanMX-GAL1pr-FLO8-HIS3 rco1Δ0::kanMX* | (Cheung et al. 2008) |
| YLH510 | L1106 | Mat a | *his3Δ200 ura3Δ0 kanMX-GAL1pr-FLO8-HIS3 rco1Δ0::kanMX sas3::URA3* |  |

**Table S3:**
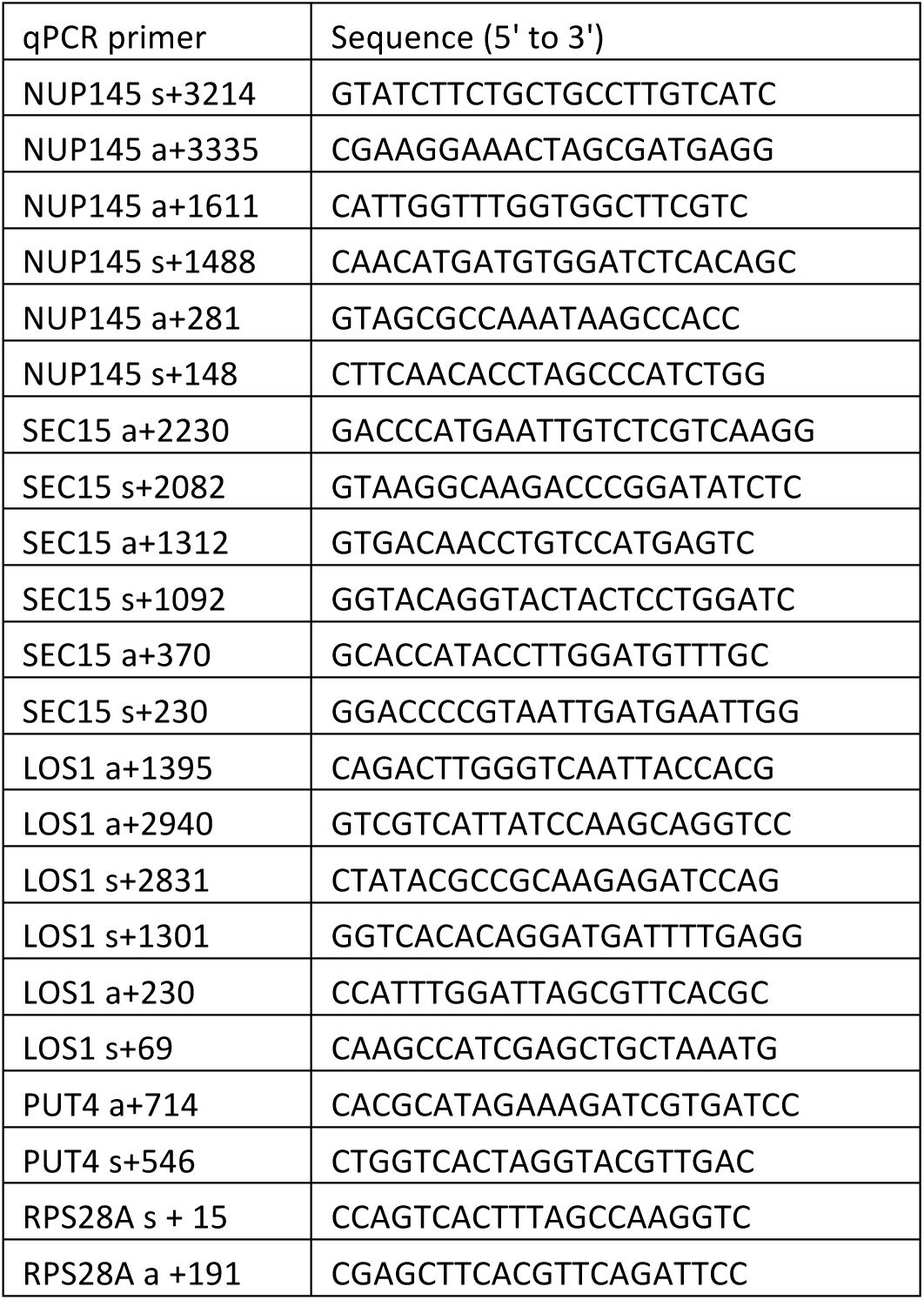
ChIP-qPCR primers

**Table S4:** Nucleosome Spearman correlation coefficients.

